# The mouse papillomavirus epigenetic signature is characterised by DNA hypermethylation after lesion regression

**DOI:** 10.1101/2021.04.19.440429

**Authors:** Allison M. Tschirley, Peter A. Stockwell, Euan J. Rodger, Oliver Eltherington, Ian M. Morison, Neil Christensen, Aniruddha Chatterjee, Merilyn Hibma

## Abstract

The β genus of human papillomaviruses (HPVs) infect cutaneous epidermis. They contribute to the development of cutaneous squamous cell carcinoma (cSCC) in individuals with epidermodysplasia verruciformis, and increase susceptibility to UV-induced cSCC. This has been demonstrated in UV-exposed mice previously infected with mouse papillomavirus (MmuPV1). However, the mechanism by which β-HPVs contribute to cSCC is unclear. We propose that viral infection leaves a DNA methylation signature following resolution of the active lesion that may contribute to increased susceptibility to UV-induced cSCC.

To test this, we carried out Reduced Representation Bisulphite Sequencing on DNA from tail skin of mice with actively infected lesions, MmuPV1-infected then healed lesions (regressed infection), and mock-infected control mice. Genome-scale DNA methylation libraries were generated and analysed for differentially methylated regions throughout the genome, and for HPV sequences.

We found that DNA of active lesions was not differentially methylated compared to matched control mice. In contrast, 834 differentially methylated fragments were identified in regressed lesions compared to mock-infected control skin. An analysis of MmuPV1 viral DNA demonstrated retention of viral DNA in some of the lesions that had regressed. Overall, the viral sequences identified showed over-representation of sequences from the E4 region. The DNA hypermethylation that we found in regressed MmuPV1 lesions may be a factor in the increased susceptibility of mice to UV-induced cSCC.

**AUTHOR SUMMARY:** Papillomavirus infections can be asymptomatic, can cause warts, and in some cases can lead to cancer. There is direct evidence for mouse papillomavirus infection resulting in increased susceptibility to UV-induced cutaneous squamous cell carcinoma in a mouse model. We propose that DNA methylation following viral infection may contribute to the increased susceptibility. We describe the DNA methylation landscape during an active infection with mouse papillomavirus and following regression of the lesion. We found that there were very few differentially methylated DNA fragments during active infection. In contrast, over 800 differentially methylated DNA fragments were identified following regression of the lesion. This is the first description of the genome-wide DNA methylation landscape for mouse papillomavirus, to our knowledge. The dramatic DNA hypermethylation that we observe following resolution of infection may contribute to a ‘hit and run’ mechanism for the increased susceptibility to UV-induced cancer by papillomaviruses.

## INTRODUCTION

Cutaneous squamous cell carcinoma (cSCC) is one of the most common cancers world-wide (1). Mortality is at least ten-fold higher in immunosuppressed individuals compared with those who are immune competent (2). Although ultra-violet (UV) light is the primary initiator of oncogenesis for cSCC, the β-human papillomavirus (β-HPV) genus has been implicated in cSCC causality. However, this proposition is confounded by the observation that, unlike cervical cancer, β-HPV DNA is not always detected in the resultant tumour. This has led to the formulation of a viral ‘hit and run’ hypothesis, whereby the β-HPV contributes to the process of carcinogenesis but is then lost from the cSCC (3).

β-HPVs are almost ubiquitously present on the skin of individuals. Unlike the mucosal family of α-HPVs, some members of which have been shown to directly cause a variety of anogenital cancers, β-HPVs are directly linked to the causality of skin malignancies only in special cases, particular systemic epidermodysplasia verruciformis (EV) (4). Several lines of *in vitro* and *in vivo* evidence have implicated β-HPVs in the development of UV-induced cSCC (5-8). For example, EV patients are more likely to develop SCC (4) and keratinocytes exposed to β-HPV E6 proteins are protected from the apoptotic pathway triggered by UVB irradiation (9).

Until recently, *in vivo* experimentation to assess the role of papillomavirus in UV-induced cSCC has been hindered by a lack of naturally occurring, accessible murine papillomavirus models. Mouse papillomavirus type 1 (MmuPV1) has more recently been described by Ingle and colleagues in Mumbai, India (10, 11). This virus is a murine homologue of the HPVs, bearing particular resemblance to the β-HPV genus in several ways including causing the formation of cutaneous papillomas following infection of immunosuppressed mice.

Immunosuppression during MmuPV1 infection leads to an increased risk of cSCC from chemical carcinogens or UV light (12, 13). It is proposed that the virus induces epigenetic changes within the host that increase the risk of UV carcinogenesis (8). It is also proposed that immunosuppression suppresses antigen-specific immune defence involving the viral E7 protein that is a CD8 T-cell target in MmuPV1 infected cells (14, 15).

The circular DNA of MmuPV1 encodes seven open reading frames (ORFs) found in other HPV types (E1, E2, E4, E6, E7, L1 and L2). Each of these ORFs is differentially expressed in the viral life cycle, similarly to other papillomaviruses (11). It has not yet been established whether all of the functions of the MmuPV1 viral proteins function the same way as HPVs, but importantly it appears that E6 and E7 are still required for oncogenesis. Similar to the cutaneous HPV types in the β family, MmuPV1 does not contain an E5 ORF (16). The E4 protein is known to accumulate at high levels during productive infection (17). The loss of E4 expression in β-HPV lesions correlates with an increase in cervical intraepithelial neoplasia (CIN) grade in cervical SCC patients (18).

Methylation of DNA is an epigenetic change that can alter expression of genes that has been linked to many cancers including cSCC (19-21). Several DNA tumour viruses such as EBV, HBV and HPV have been found to dysregulate host gene DNA methylation (22). α-HPVs regulate methylation (23), however the effects of this methylation on host susceptibility to cancer once the virus infection has resolved is unknown. Methylation occurs predominantly at CpG dinucleotides, where a cytosine is followed by a guanine. CpG dinucleotides are generally underrepresented in the genomes of both DNA viruses and their hosts (24).

Cellular gene methylation in the context of high-risk α-HPVs or cSCC is well-documented (19, 25-27). Cancers associated with HPV are similar to most other cancers in their propensity towards global hypomethylation (28). Some of the most frequently hypermethylated genes in cSCC include *CADM1, CDH1* and *DAPK1* (19). Methylation of *DAPK1* is associated with cSCC development and progression.

There is a paucity of data on DNA methylation in the context of MmuPV1 infections, and this study is the first to describe DNA methylation in resolved lesions following MmuPV1 infection.

## RESULTS

### Immune-suppressed BALB/c mice develop and maintain active MmuPV1 infection

BALB/c mice demonstrate intermediate susceptibility to MmuPV1 infection when immunosuppressed with CsA (15). Handisurya *et al*. (15) found that five out of eight (63 %) BALB/c mice produced MmuPV1 lesions following infection. In this study, the proportion of mice that were infected was much lower (20 %), and some mice required a second dose of virus for a visible lesion to appear. Lesions visible to the naked eye appeared on the tail of CsA treated mice between 49 and 63 days post infection, and any mice in the infected groups that did not develop visible lesions were excluded from the study (Fig. 1A). The mean lesion length on the infected mice was 2.4 ± 0.3 mm. CsA treatment was withdrawn for the regressor group to promote lesion regression. All mice in the regressed group had healed their lesions (assessed by visual and tactile tests) by 31 days post CsA withdrawal.

**Fig. 1.**
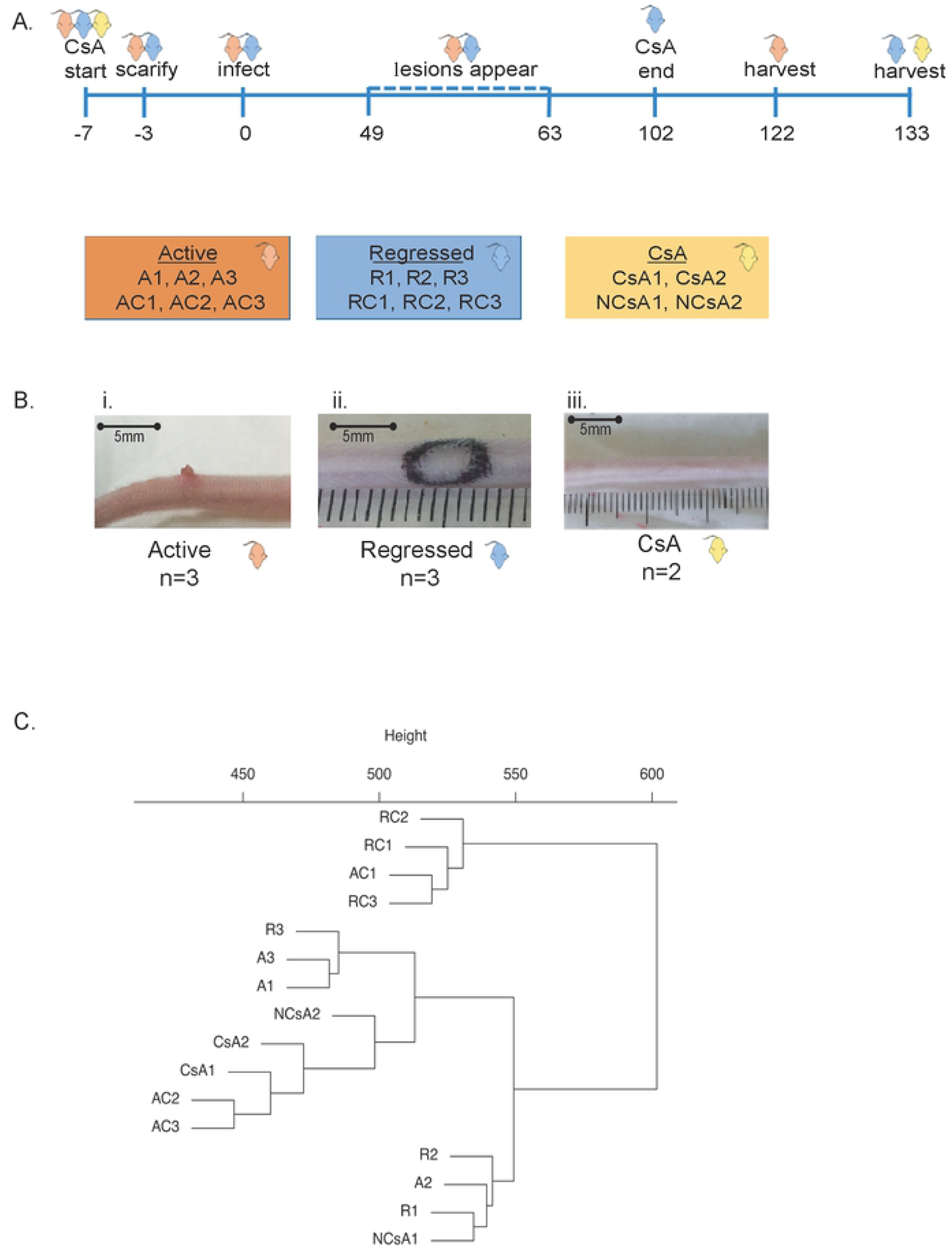
Experimental setup for mouse model and preliminary methylation analysis. (A) Timeline of infection model. Colour-coded mouse images represent the three groups as shown in the boxes below. Boxes show sample names of each group and their associated control samples. (B) Representative images of the experimental groups show mouse tails at sample location; (i) Active, (ii) Regressed (original lesion location delineated in black), (iii) CsA. (C) Hierarchical clustering dendrogram of all high-coverage autosomal fragments for the 16 RRBS libraries generated.

Reduced Representation Bisulphite Sequencing (RRBS) was carried out on tissue samples from treated and control mice, in order to determine the effects of viral infection and regression on the methylation profile, and CsA treatment and scarification. On average, we generated 9.3 x 10^6^ ± 0.5 x 10^6^ uniquely aligned reads for each of the samples tested (Table 1). The mean mapping efficiency was 47 % ± 1.3 %, which is consistent with the expected level of efficiency for a bisulphite converted genome.

**Table 1.** Methylation alignment and efficiency by individual sample and number of differentially methylated fragments (DMFs) using Benjamini-Hochberg correction for each group comparison.

| Comparison | Individual | No. of unique alignments | Mapping efficiency | No. of DMFs at 5% FDR |
| --- | --- | --- | --- | --- |
| Regressed | R1 | 6,942,498 | 44.3% | 834 |
|  | R2 | 7,861,430 | 48.6% |  |
|  | R3 | 7,240,969 | 47.0% |  |
| Reg. Control | RC1 | 11,964,235 | 50.6% |  |
|  | RC2 | 12,197,807 | 49.8% |  |
|  | RC3 | 9,824,427 | 39.5% |  |
| Active | A1 | 7,449,228 | 49.6% | 0 |
|  | A2 | 10,712,796 | 33.7% |  |
|  | A3 | 6,442,790 | 49.5% |  |
| Act. Control | AC1 | 13,217,138 | 38.3% |  |
|  | AC2 | 9,198,760 | 49.8% |  |
|  | AC3 | 10,374,183 | 51.9% |  |
| No CsA | NCsA1 | 6,434,780 | 47.8% | 0 |
|  | NCsA2 | 6,853,519 | 49.1% |  |
| CsA | CsA1 | 9,428,602 | 49.5% |  |
|  | CsA2 | 12,426,447 | 52.7% |  |
| Non-scarified | CsA1, CsA2 |  |  | 0 |
| Scarified | AC1, AC2, AC3 |  |  |  |
| Active | A1, A2, A3 |  |  | 0 |
| Regressed | R1, R2, R3 |  |  |  |

When we compared tissues from the regressed mice and their matched controls we found that there was a marked difference in DNA methylation between the groups. The comparison of the regressed (R1 – 3) versus the regressed control group (RC1 – 3) yielded 834 significantly differentially methylated fragments (DMFs) with a False Discovery Rate (FDR) of 5 % or less using a stringent multiple test correction (Benjamini-Hochberg (Table 1).

In contrast, there was little difference in DNA methylation between the tissues from actively infected mice and their controls. Only five DMFs were identified in the actively infected group (A1 – 3) when compared to the matched control samples (AC1 – 3). Treatment of mice with CsA alone (CsA1 – 2 v. NCsA1 – 2) or scarification alone (A1 – 3 v. CsA1 – 2) had little effect on differential methylation between the condition and the control.

To assess the relationship of global RRBS methylation between the samples that were tested, we performed hierarchical clustering of all of the analysed high coverage autosomal fragments from the 16 RRBS methylomes in this study (Fig. 1C). We found that the infected samples grouped into two clusters, irrespective of whether they were actively infected or if the lesion had regressed.

Samples from uninfected mice generally grouped into two other clusters, separated from the infected mice.

### Genome-scale DNA methylation identifies extensive hypermethylation in regressed skin

We carried out further analysis to determine if the DMFs were hyper-or hypomethylated. The striking finding from this analysis was that 98% of the DMFs identified in the regressed group were hypermethylated (Mann-Whitney U, *P* < 0.0001) (Fig. 2A).

**Fig. 2.**
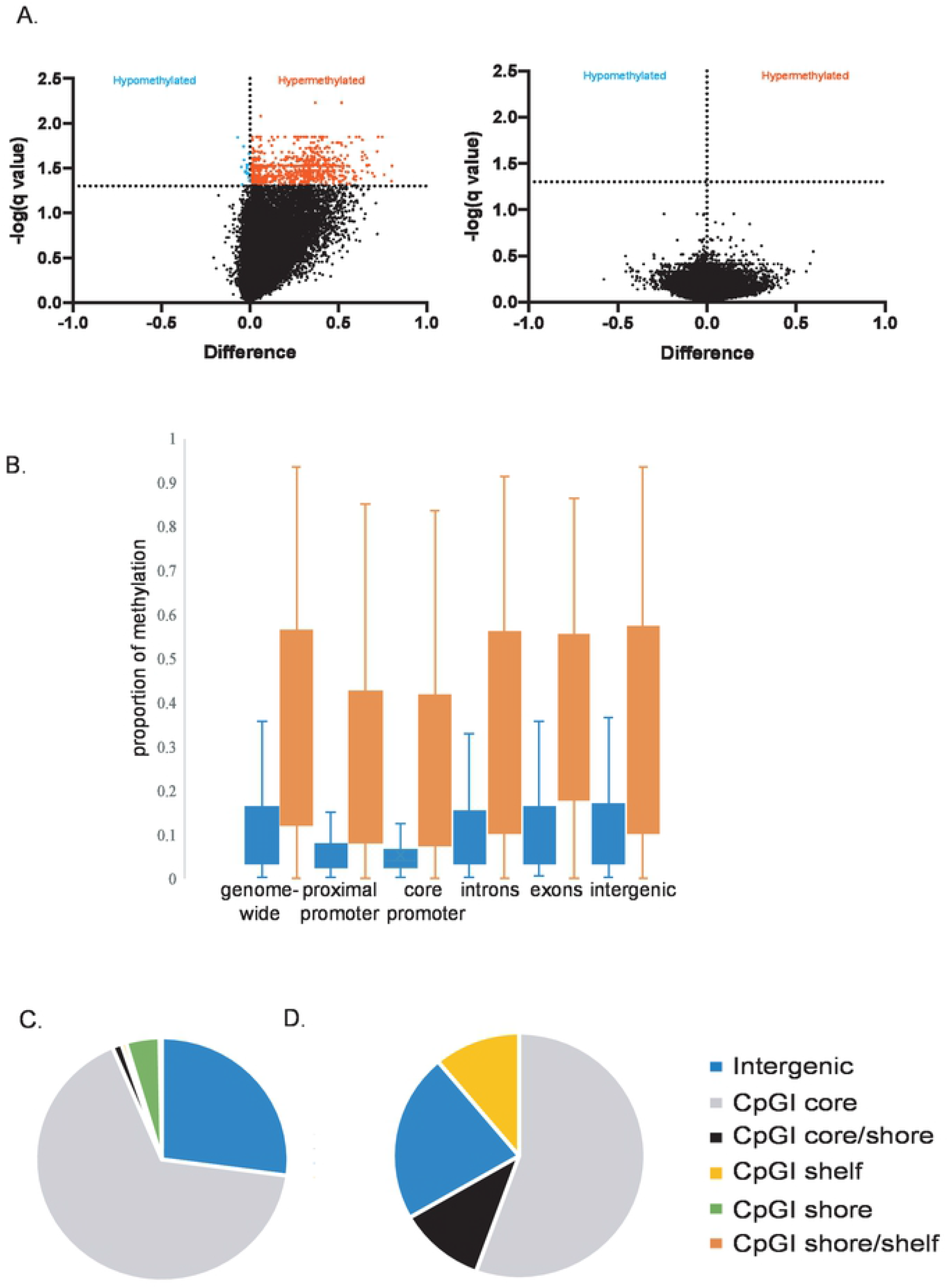
Global methylation description. (A) Difference plots of differentially methylated fragments (DMFs) in comparison of regressed to regressed control tissue (left) and active to active control tissue (right). Horizontal dotted line represents significance cut-off of 5 % FDR. (B) Proportion of methylated DMFs by genomic region for regressed (blue) and regressed control tissues (red). Data are represented in quartiles with whiskers indicating outliers outside of upper and lower quartiles. (C) Within CpG islands, differentially hypermethylated regions (n = 328) and (D) hypomethylated regions (n = 9) were located predominantly in the core.

To further probe the DMFs in the regressed MmuPV1 lesions relative to their location in the genome, we generated DNA methylation maps for the specific genomic regions. We found that methylation was increased overall in the DMFs from all genomic regions that we assessed in regressed skin when compared with controls (Fig. 2B). Significantly increased methylation was detected in DMFs genome-wide (Mann-Whitney U, *P* < 0.0001), proximal to gene promoters (defined as +/-1 kb of transcription start site) (Mann-Whitney U, *P* < 0.0001), in the core promoter region (defined as +/-500 bp of transcription start site) (Mann-Whitney U, *P* < 0.0001), in gene introns (Mann-Whitney U, *P* < 0.0001) and gene exons (Mann-Whitney U, *P* < 0.0001), and in intergenic regions (Mann-Whitney U, *P* < 0.0001).

To further investigate the methylation landscape, we performed an analysis of the wider CpG island topography (10 kb upstream to 1 kb downstream of transcription start sites). CpG islands are defined as stretches of DNA 500–1500 bp long with a CG:GC ratio of more than 0.6 (29). We found that most of the hypermethylation was in the CpG island cores. Of 328 hypermethylated fragments that were detected within the region of interest, 66 % were found in the core, 4 % were in the CpG island shore (up to 2 kb upstream and 2 kb downstream of the CpG island core), 1 % were in the interface between core and shore (termed core-shore) and <1 % were in the shelf (up to 2 kb upstream and 2 kb downstream of the shore). 27 % of the regions were outside of these CpG island features (Fig. 2C). Of the nine hypomethylated fragments that were detected, five were in the CpG island core (Fig. 2D).

### Genes in the promoter region are skewed extensively towards hypermethylation in regressed skin

As promoter DNA methylation strongly influences expression from the corresponding gene, we specifically investigated differential methylation at the core promoter regions. Of the 221 DMFs in these regions, 214 were hypermethylated and seven were hypomethylated. When comparing individual samples within the groups, all genes with > 20 % methylation difference were hypermethylated in the regressed group (Fig. 3A). This was also the case at > 40 % methylation difference (Fig. 3B), where all genes were hypermethylated in regressed tissue compared to control tissue.

**Fig. 3.**
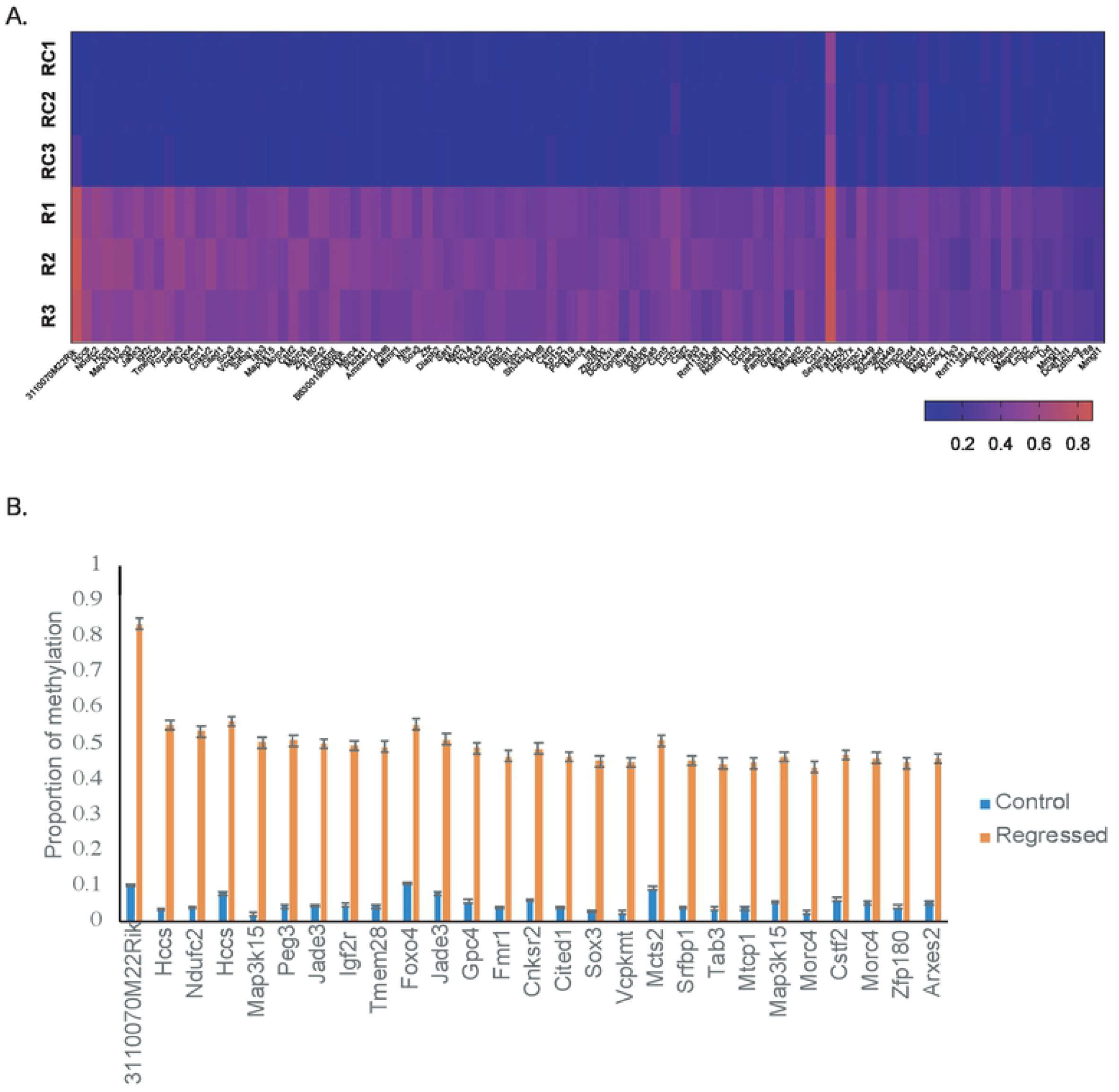
Extensive hypermethylation identified in regressed skin. (A) DMFs in the core promoter region with > 20% methylation difference between regressed and control groups by individual sample. (B) DMFs in the core promoter region with > 40% methylation difference between regressed and control groups. Error bars display standard error of the mean.

### Hypermethylated genes in regressed skin are enriched for cellular senescence and cancer related pathways, and for CTCF and RNA Polymerase II binding

To document the function of the genes that were hypermethylated in the core promoter regions of regressed skin, we performed functional enrichment analysis using two different sources (KEGG and Wiki Pathways for Mouse). We found genes involved in cellular senescence/cell cycle, mRNA processing, and p53 signalling were significantly enriched (*P* < 0.05) in both the analyses (Fig. 4A). Disease associated pathways, particularly cancer pathways, were also enriched in our analysis. To place our findings in a broader epigenomic context, we utilized histone modification ChIP-Seq data from the ENCODE project as well as from the Epigenomics roadmap. The genes harbouring hypermethylation in the core promoters of regressed skin were predominantly enriched for the repressive chromatin mark H3K9me3 (in several cell types, see Fig. 4B). We also found significant enrichment for several active histone marks, especially H3K27ac and H3K29ac (Fig. 4B). Overlap analysis of ENCODE transcription factor ChIP-seq data identified enrichment of several transcription factors for the hypermethylated genes (Fig. 4C). The most enriched (significant in many cell types) transcription factors were CTCF and POLR2A (RNA Polymerase II Subunit) (Fig. 4C).

**Fig. 4.**
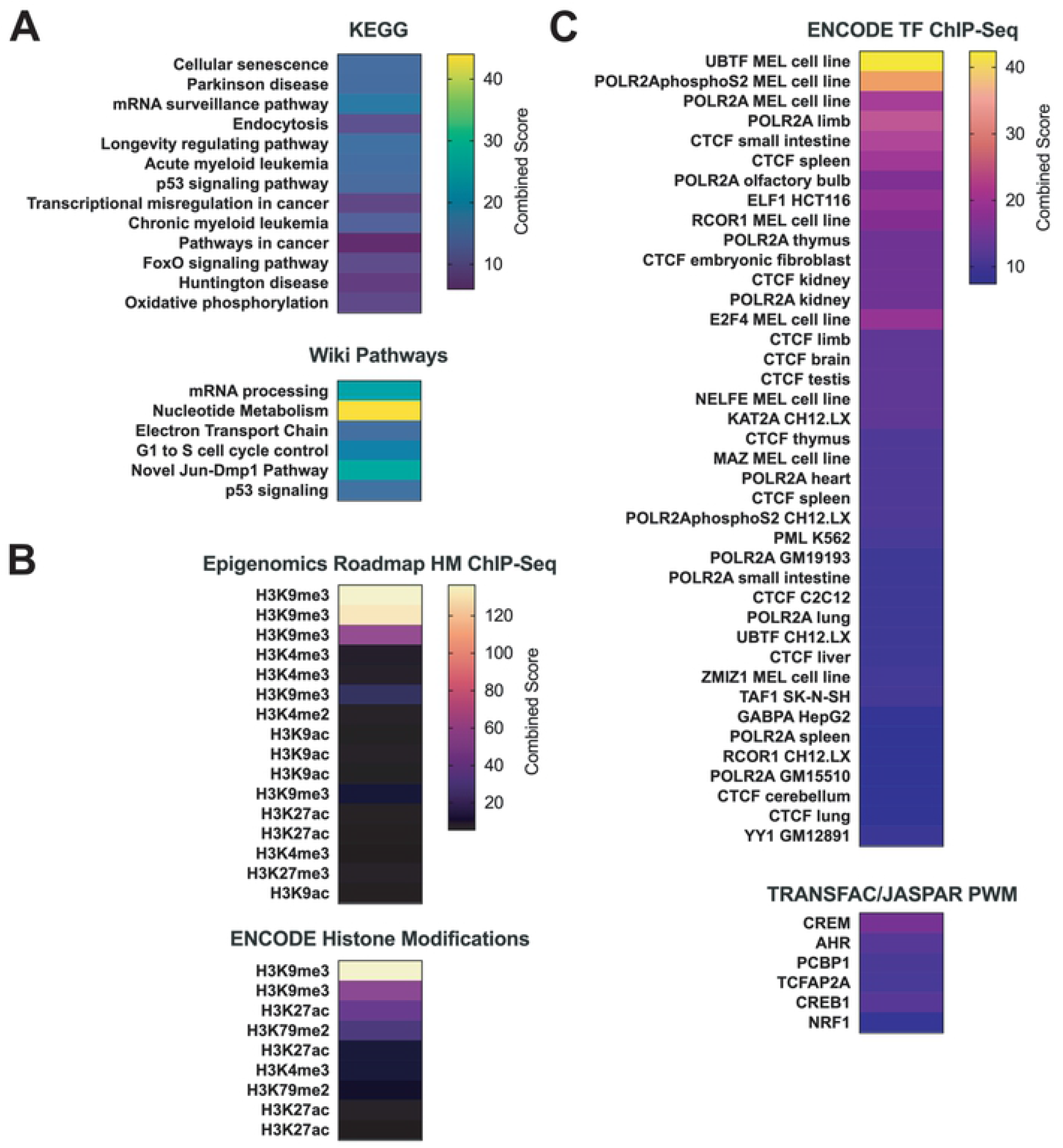
Pathway and transcriptional function analysis of differentially methylated fragments (DMFs). Enrichr analysis with default parameters was used to identify pathways, histone modifications and transcription factor targets associated with DMFs in core gene promoter regions. Individual heatmaps show: (A) enriched KEGG and Wiki Pathways (*P* < 0.05); (B) enriched histone modifications from Epigenomics Roadmap and ENCODE datasets (*P* < 0.01); and (C) Transcription factor targets from ENCODE ChIP-Seq and TRANSFAC/JASPAR positional weight matrices (*P* < 0.01).

### MmuPV1 viral reads are highly concentrated in the E4 portion of the genome

We probed the data for MmuPV1 viral reads to determine whether viral DNA remained after lesion regression (Fig. 5A). Traces of the viral genome were present in all actively infected tissue (5.7 %, 0.2 % and 5.4 % of viral reads were viral in origin for samples A1 to A3, respectively). Interestingly, viral reads were also detected in two of three regressed tissues (7.2 %, 0.0 % and 5.3 % for samples R1 to R3, respectively). The small number of background reads mapping to the MmuPV1 genome in uninfected groups most likely occurred through the misalignment of bisulphite reads, as previously reported (30) (S1 Table). The most intriguing finding of viral read analysis was the high proportion of viral reads within the E4 region of the MmuPV1 genome (Fig. 5B, C). In two of the actively infected samples (A1 and A3), more than half of all reads aligned to this region. In one of the regressed lesions, sample R1, 74 % of the viral reads mapped to E4. The L2 region also had high clustering, contributing up to 25 % of all viral reads in the actively infected samples.

**Fig. 5.**
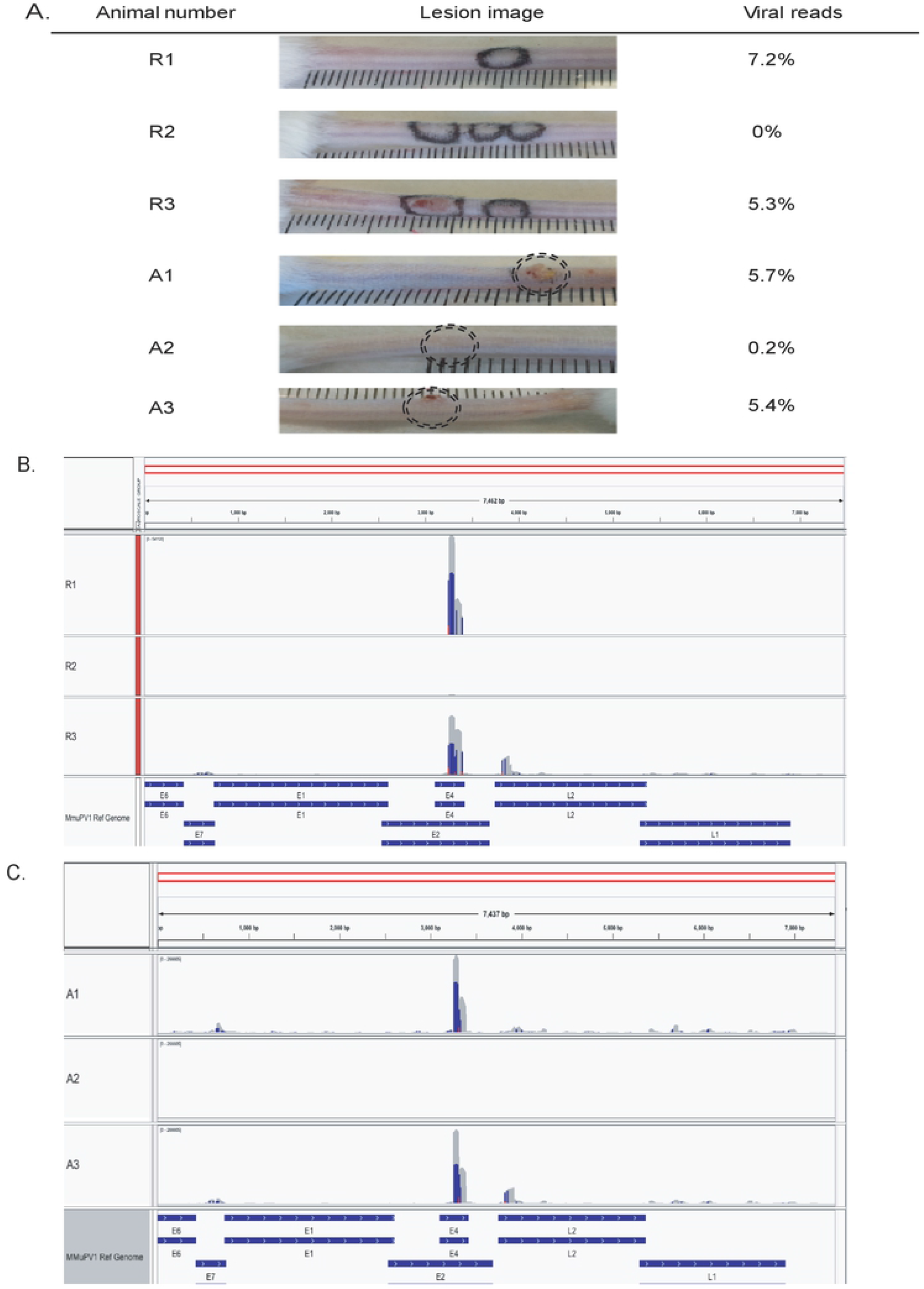
MmuPV1 viral reads within individual samples. (A) Images at time of harvest. Marker pen was used to delineate the area of the lesions at the height of infection (B-C). Coverage and distribution of viral reads visualised using IGV software. Coverage depth of reads set to group auto-scale for (B) regressed (0-266,605) and (C) active groups (0-541,120).

## DISCUSSION

Here we show the first reported analysis of DNA methylation following regression of MmuPV1 infection. Interestingly, differences in methylation were only detected following lesion regression, and not during active infection. These differences were highly skewed towards hypermethylation.

In our analysis, *DAPK1* was hypermethylated in core promoters in regressed skin compared to control tissue. Hypermethylation is associated with cSCC as reported by Li and colleagues, who found that methylation of *DAPK1* was associated with cSCC but in their case it did not correlate with the presence of cutaneous HPV (19). *DAPK1* is considered likely to be a tumour suppressor. Our finding is supported by RNAseq analysis performed by Hu and colleagues on MmuPV1 infected tissue where *DAPK1* was downregulated by 0.76 log_2_ fold change from infected (with no lesion) to infected (with lesion) (31). The hypermethylation of the tumour suppressor gene *DAPK1* indicates that this gene is ‘switched off’ in MmuPV1 regressed lesions. *DAPK1* presence in the core promoter region suggests that its hypermethylation may influence the *DAPK1* expression profile, and thus could be a possible route to increased susceptibility to cSCC after regression of HPV infection.

The identification of hypermethylation of POLR2A is consistent with the observation that one of the enriched pathways that we identified was mRNA processing, suggesting a role of hypermethylation in RNA processing in regressed skin. We also identified hypermethylation of the CTCF promoter. Variable CTCF binding has been strongly linked with differential DNA methylation in multiple human cell types and it has been shown that CTCF binding patterns are markedly different in normal versus immortal human cells. In immortal cells, disruption of CTCF binding was strongly associated with hypermethylation (32). Our analysis provides evidence for hypermethylation and disruption of CTCF binding in regressed skin compared to normal skin controls.

Interestingly, *MSH6* and *PAPD7* are DNA mismatch repair genes that were also hypermethylated within the core promoter region of regressed skin (5 % and 6.5 % methylation difference, respectively). *MSH6* is important for UVB-induced apoptosis (33, 34). Hypermethylation of *MSH6* in regressed tissue prevents skin cells from effectively undergoing UVB-induced apoptosis. *PAPD7* has been found to be upregulated by HSV1 infection (35). Its hypermethylation indicates that the damage response no longer occurs. Indeed, Uberoi *et al*. (13) found that immune-competent mice infected with MmuPV1 and exposed to UV radiation develop lesions that progress to cSCC. Key genes such as *MSH6*, whose promoters are hypermethylated by MmuPV1, may render the skin more sensitive to the damaging effects of UV, providing the virally mediated susceptibility ‘hit’ that promotes UV-induced SCC.

Our analysis of viral reads found evidence of viral DNA in regressed tissues with no remaining visible lesion. To our knowledge, this is the first evidence of the presence of MmuPV1 DNA in infected tissue where lesions were no longer observable. Viral gene expression has been detected by Xue *et al*. (17) well before lesions are observed. These investigators also observed asynchronous papillomavirus growth, which was similarly found by our analysis of viral reads, where one visible lesion in the actively infected group (sample A2) yielded only 0.2% viral reads while skin with no visible lesion in the regressed group (sample R1) showed the highest level of viral DNA at 7.2 %.

The low viral reads of sample A2 could be partly explained by the size of the lesion, which was smallest of the three active lesions (height: 0 mm, length: 2 mm, width: 1 mm) and could contribute to its methylation profile clustering with two of the samples from the regressed lesion group (Fig. 1C). Interestingly, R1 which had the highest level of viral DNA clustered with A2 (0.2 %), R2 (0 %) and NCsA1 (did not receive virus). Therefore, it is not solely the amount of viral DNA that dictates the methylation patterns of the samples, nor is it the current or past immune suppression, as NCsA1 did not receive cyclosporin at any stage. Rabbit oral papillomavirus DNA and RNA has been found to persist for at least a year post lesion regression (36). The high viral reads in two out of three regressed lesions suggests viral latency in MmuPV1 is also possible. The diversity in the quantity of viral DNA found could be explained by the stage of epithelial differentiation that the cells were in when collected. A progression of this research could involve re-establishment of immunosuppression in regressed animals to see if the latent infection re-emerges at the site of the original lesion.

The most striking finding within the analysis of viral reads was the high clustering of reads in the E4 region. E4 protein is found to be abundant in upper layers of the epithelium of productive lesions (37). E4 expression is followed by the late structural proteins, explaining the concurrent, though considerably lower, expression of L2. This pattern was also found by Xue *et al*. (17), who looked solely at active MmuPV1 infections in the tail, muzzle and ear of mice.

There is considerable variability in the pattern of expression of different papillomavirus types, depending on the animal model and tissue tropism (38). High levels of E4 transcripts in the upper layers of the epithelium, similarly to the DNA viral reads found here, follow the pattern found in low-grade squamous intraepithelial lesions of high-risk cervical HPV infections (39). In an analysis of 92 different HPV genomes, the E4 region was found to have the highest proportion of CpG sites (40). This is despite the fact that CpG sites tend to be under-represented in viruses, which may be a way to avoid methylation by host methyltransferases or CpG mediated immune responses.

In summary, our study outlined the methylation landscape and viral content and distribution of MmuPV1 in regressed lesion tissue. We highlight several genes hypermethylated in the core promoter regions of regressed skin that could influence susceptibility to cSCC without the need for active lesions to be present. In future it would be useful to leave regressed skin for longer and continue to monitor viral content. Experiments using UV as a co-factor would also aid in validating the function or loss of function of the identified genes through methylation.

## MATERIALS AND METHODS

### Animals

Female specific pathogen-free (SPF) BALB/c mice, aged 8 - 12 weeks, were obtained from the Hercus Taieri Research Unit at the University of Otago and housed in a SPF environment. A total of 63 mice were initially in the study. For systemic immune suppression, Cyclosporin A (CsA, Novartis Neoral®, 100 mg/mL) diluted in PBS was administered to the 30 control and 30 infected mice by gavage five times per week (75 mg/kg, 11.25 mg/mL) starting one week prior to MmuPV1 infection. The remaining three mice were the non-CsA treated mice. Thirty of the mice were infected with MmuPV1 and were monitored for the development of visible lesions and the other thirty mice were mock infected with PBS. Mice that had not developed visible lesions were re-infected at day 49. Six animals developed visible lesions, and the remainder of the infected mice that did not develop visible lesions were excluded from the study. The final groups (Fig. 1) consisted firstly of mice (n = 3) that were harvested at day 122 with “active” lesions (A1, A2, A3) and three matched mock-infected controls (AC1, AC2, AC3), and secondly, mice that had active lesions but were harvested at day 133 with “regressed” lesions (R1, R2, R3) following the withdrawal of CsA at day 102, and three matched CsA treated, mock-infected controls (RC1, RC2, RC3). Groups of two CsA treated mice (CsA1, CsA2) and two controls (NCsA1, NCsA2) were included in the study to test whether any changes in DNA methylation occurred as a result of CsA treatment alone. Commencing at day -3, Baytril was added to drinking water for all immune suppressed mice to reduce the risk of bacterial infection.

### Ethics statement

All animal experiments were approved by the Animal Ethics Committee at the University of Otago (ethical approval AEC56/14) and were performed in accordance with the New Zealand Animal Welfare Act (1999) and adhered to National Animal Ethics Committee (Ministry of Primary Industries, NZ) guidelines.

### MmuPV1 infection

Mouse papillomavirus was isolated from lesions on the tails of Foxn1^nu^/Foxn1^nu^ immunocompromised mice and diluted 1:10 in PBS (41). On day -3, tail skin was scarified using a hand-held rotary tool with a felt wheel attached (42). Animals were anaethetised with Isoflurane, and a 1.5 cm long mark was placed at the base of the tail using a felt tip pen, and the rotary tool was moved 10 times along the mark at a speed of approximately 15,000 rpm to remove the upper epidermal layer. Marcain was applied dropwise over the wound area for pain relief. At day 0, mice were placed in restrainers and the scarified region was scored lightly with a needle tip. Inoculation was performed by placing 20 μL of prepared virus onto the scored region of the scarified area (Fig. 1A). Mock-infected mice received PBS. At day 49, animals who had yet to show signs of a lesion were re-infected with 20μl of prepared virus. At day 102, CsA was discontinued for the regressor group, to allow the lesions to heal naturally.

### Sample preparation

Beginning at day 102 for the regressor group, the lesions were outlined with marker pen for identification of tissue to be harvested once lesions regressed. The actively infected group mice were culled at day 122 post-infection. Tail skin was removed and epidermal dissociation of the epithelium was performed following the protocol of Lichti *et al*. (43). DNA was extracted from epidermal tissue (approximately 3 - 5 mm^2^ per lesion) using a DNAeasy blood and tissue kit (Qiagen). Regressed and CsA group mice were culled at day 133 post-infection, once lesions were no longer palpable, and the regressed group was culled at 31 days post withdrawal of CsA.

### RRBS library preparation

RRBS was performed on 16 samples of extracted genomic DNA as previously described (44-46). Briefly, DNA was digested into fragments using the MspI enzyme, followed by end-repair and ligation of sequencing adaptors. The fragments were size selected (150 - 325 bp) and bisulphite converted prior to a PCR amplification step. The quality of the 16 libraries was confirmed using bioanalyser traces before being sequenced on an Illumina HiSeq2500 machine (Table 1).

### DNA methylation analysis

Differential methylation analysis was performed using the previously reported differential methylation analysis package (DMAP) (47). Briefly, only fragments with at least two CpG sites and at least ten hits per CpG were included in the data (hereon called high coverage fragments). We have previously described this fragment-based approach (48-51). High coverage fragments were mapped to the mouse genome and the closest protein coding genes were annotated. An ANOVA test was applied to the fragments of the experimental and control groups and regions showing methylation differences with a fold change of ≥ 1.5 and significant *P*-values were identified. Five different group comparisons were performed (Regressed vs. Control, Active vs. Control, Active vs. Regressed, CsA vs. no CsA, Scarified vs. Non-scarified) (Fig. 1B). Active and regressed tissues were the experimental groups. Non-scarified vs. scarified was a methylation comparison only and compared animal samples from the CsA and actively infected control groups. These control tissues were tested to determine whether scarification impacted on methylation. CsA vs. no CsA tissues was assessed to determine whether CsA administration impacted methylation.

### Analysis of viral reads and pathway analysis

Methylation data were aligned to the MmuPV1 genome using bowtie in the Bismark Bisulphite Read Mapper (52). Sequences that mapped to the MmuPV1 genome were also aligned to the *Mus musculus* genome to ascertain whether mapping was unique to the viral genome. The mapper BSMAP(z) was then used to verify the alignments shown by Bismark. This mapping software confirmed that the reads were not indicating artefactual behaviour of Bismark alone. The Integrative Genomics Viewer (IGV) was used to visualise reads within the MmuPV1 genome (53). Lanes were auto-scaled to show proportionate representation between the samples.

Unsupervised hierarchical clustering was performed in the R environment using the Euclidean distance metric of all analysed high coverage autosomal fragments (n = 126,681) from the 16 RRBS methylomes. Pathway analysis was performed on a gene list containing the 221 genes differentially expressed in the core promoter region using the online platform Enrichr (54). The enrichment *P* values calculated by Enrichr are from a modified Fisher’s exact test, which is a proportion test that assumes a binomial distribution and independence for probability of any gene belonging to any set.

### Statistical analyses

All data are presented as the mean ± standard error of the mean and statistical analyses were done in GraphPad Prism7®. Mann-Whitney U was used to test for differences between groups. The original FDR method of Benjamini and Hochberg under the multiple T tests function was used to increase the power of the data and identify significant *P* values with a false discovery rate (FDR) of 5%.

### Data availability

The RRBS data generated as part of this study were submitted to Gene Expression Omnibus (GEO) with the accession number GSE160868.

## AUTHOR CONTRIBUTIONS

Conceptualisation of research: MH, AT, AC, NC, IM, OE; Data Curation: AT, PS, AC; Formal Analysis: PS, AT, MH, AC, ER; Methodology: AT, NC, MH, AJ; Software: AC, PS, ER; Funding acquisition: MH, AT, AC, IM; Writing: AT, OE, MH, AC, IM, NC.

ACKNOWLEDGMENTS

We thank Jiafen Hu (Pennsylvania State University) for supplying additional RNAseq data from MmuPV1 tissues. The authors declare no conflicts of interest. AC would like to thank the Rutherford Discovery Fellowship (Royal Society of NZ) for their support.

---

**S1 Table.** Viral reads were present within regressed lesions. Total and percent viral reads within E4 region.

| Group | Sample | Total reads | Viral reads | % viral reads | % in E4 | Infection status |
| --- | --- | --- | --- | --- | --- | --- |
| Regressed | R1 | 15,674,849 | 1,129,427 | 7.2 | 74 | Regressed lesion |
|  | R2 | 16,166,560 | 5,871 | 0 | 0 | Regressed lesion |
|  | R3 | 15,392,780 | 819,114 | 5.3 | 59.9 | Regressed lesion |
| Reg. Control | RC1 | 23,666,799 | 3,713 | 0 | 0 | Mock-infected |
|  | RC2 | 24,496,535 | 2,868 | 0 | 0 | Mock-infected |
|  | RC3 | 24,877,419 | 4,143 | 0 | 0 | Mock-infected |
| Active | A1 | 15,020,374 | 849,677 | 5.7 | 52.1 | Active lesion |
|  | A2 | 31,835,719 | 48,521 | 0.2 | 11.4 | Active lesion |
|  | A3 | 13,025,548 | 708,976 | 5.4 | 56 | Active lesion |
| Act. Control | AC1 | 34,512,945 | 6,246 | 0 | 0 | Mock-infected |
|  | AC2 | 18,479,427 | 2,534 | 0 | 0 | Mock-infected |
|  | AC3 | 19,996,210 | 2,475 | 0 | 0 | Mock-infected |
| CsA | CsA1 | 19,044,226 | 1,271 | 0 | 0 | CsA/No infection |
|  | CsA2 | 23,584,967 | 1,829 | 0 | 0 | CsA/No infection |
| No CsA | NCsA1 | 13,470,444 | 1,106 | 0 | 0 | No CsA/Not inf. |
|  | NCsA2 | 13,946,497 | 1,282 | 0 | 0 | No CsA/Not inf. |

